# Macrophages inhibit and enhance endometriosis depending on their origin

**DOI:** 10.1101/2020.04.30.070003

**Authors:** Chloe Hogg, Priya Dhami, Matthew Rosser, Matthias Mack, Daniel Soong, Jeffrey W Pollard, Stephen J Jenkins, Andrew W Horne, Erin Greaves

## Abstract

Macrophages are intimately involved in the pathophysiology of endometriosis, a chronic inflammatory disorder characterized by the growth of endometrial-like tissue (lesions) outside the uterus. By combining genetic and pharmacological monocyte and macrophage depletion strategies we determined the ontogeny and function of macrophages in a mouse model of induced endometriosis. We demonstrate that lesion-resident macrophages are derived from eutopic endometrial tissue, infiltrating large peritoneal macrophages (LpM) and monocytes. Furthermore, we found endometriosis to trigger continuous recruitment of monocytes and expansion of CCR2+ LpM. Depletion of eutopic endometrial macrophages results in smaller endometriosis lesions, whereas constitutive inhibition of monocyte recruitment significantly reduces peritoneal macrophage populations and increased the number of lesions. We propose a putative model whereby endometrial macrophages are pro-endometriosis whilst newly-recruited monocyte-derived macrophages, possibly in LpM form, are ‘anti-endometriosis’. These observations highlight the importance of monocyte-derived macrophages in limiting disease progression.

## Introduction

Macrophages are exceptionally diverse cells present in all tissues of the body that perform functions vital for immunity, development, tissue homeostasis and repair following injury. They modify their role depending on signals received from their local microenvironment and accordingly exhibit high degrees of transcriptional and phenotypic heterogeneity and tissue-specific function^1, 2^. Macrophages differ in their ontogeny. Whilst early studies suggested macrophages were continually replaced by circulating blood monocytes, more recent lineage-tracing experiments demonstrated that most tissue-resident macrophages (exceptions include gut, dermis and heart) are derived from embryonic precursors that seed tissues prior to birth and are maintained by self-renewal or longevity^3, 4, 5^. Tissue-resident and monocyte-derived macrophages play distinct roles both in health and disease^6^. Usually, tissue-resident macrophages play tissue specific homeostatic roles as well as core functions such as clearance of dying cells. On the other-hand monocyte-derived macrophages that are recruited to tissues during inflammation secrete pro-inflammatory cytokines, help clear infection and regulate the immune response^6^. Thus, in pathological situations, macrophages are a heterogenous population. For example, during acute liver injury hepatic resident macrophages (Kupffer cells) become activated and recruit monocytes that initially promote liver injury, but then subsequently differentiate into inflammatory macrophages and help resolve injury and drive regeneration^7^. In pancreatic cancer, both tissue-resident and monocyte-derived macrophages populate the tumor and increase in density as the cancer progresses. The two populations are transcriptionally diverse and depletion studies revealed that only tissue-resident macrophages are responsible for driving tumor progression^8^. These findings also highlight that under disease-modified conditions tissue-resident macrophages can become adapted such that they promote disease.

The peritoneal cavity hosts two main macrophage populations: a predominant population expressing high levels of EGF-like module-containing mucin-like hormone receptor-like 1 (EMR1 / F4/80) and low levels of Major histocompatibility class II (MHCII) known as large peritoneal macrophages (LpM), and a less abundant population that are F4/80^lo^, MHCII^hi^ (small peritoneal macrophages; SpM). LpM are considered to be tissue resident macrophages and are largely embryonically derived, however it is now understood that they are gradually replaced by monocytes over time in a sexually dimorphic manner, that occurs more quickly in males^9, 10^. The transcription factor GATA6 is highly expressed by all LpM in response to retinoic acid and drives a significant proportion of the tissue specific transcriptional signature of these cells^11^. The SpM population is a more heterogeneous population comprising monocyte-derived macrophages and dendritic cells that are continually replenished from the blood^9, 12^. Inflammatory challenge in the peritoneal cavity can result in recruitment of large numbers of inflammatory macrophages that are transcriptionally distinct from SpM^13^, and a loss in LpM (the so-called macrophage disappearance reaction (MDR)), which results from formation of cell aggregates^14^ or cell death^15^. An exception to this is helminth infections which are characterized by expansion of LpM in response to local Th2 cytokine production^16 17, 18, 19 20^.

Endometriosis is a chronic inflammatory condition where tissue similar to the endometrium grows ectopically, usually in the peritoneal cavity as ‘lesions’^21^. The condition impacts an estimated 190 million women worldwide and is associated with debilitating pelvic pain and infertility^22^. Currently, therapeutic options are very limited, with the gold-standard treatments being surgical removal of lesions or suppression of ovarian hormones. Surgery is associated with high recurrence rates and ovarian suppression is contraceptive and often has unwanted side effects. A high abundance of macrophages is reported both in the peritoneal cavity and in lesions of women with endometriosis^23^. It is clear that macrophages are intrinsically linked with the pathophysiology of endometriosis where they enhance establishment, proliferation and vascularisation of lesions^24, 25^. They are also critical in promoting innervation of lesions and concomitant sensitization of nerve fibres, thus contributing to pain in the condition^26, 27^. Evidence from a syngeneic mouse model of induced endometriosis indicates that donor endometrial macrophages as well as host-derived macrophages can be identified in endometriosis lesions^28^. However, the exact origins of the host-derived macrophages and specific functions of the different populations remains to be determined.

Since macrophages play such a key role in many aspects of the pathophysiology of endometriosis they represent an attractive therapeutic target. However, the development of a viable immune-therapy targeting ‘disease-promoting’ macrophages or enhancing the function of ‘protective’ macrophages requires a comprehensive understanding of the origin and function of lesion-resident and associated peritoneal macrophages. In this study we have characterized the origin of endometriosis lesion-resident macrophages and examined the dynamics of peritoneal cavity macrophage populations. Finally, we have used a combination of transgenic and pharmacological approaches to selectively deplete different populations of macrophages to assess their impact on development of endometriosis lesions.

## Results

### Endometriosis lesion-resident macrophages have multiple origins

We induced endometriosis in wild-type C57BL/6 mice by injecting ‘menses’-like endometrium from MacGreen (*Csf1r-eGFP*, macrophages are GFP+)^29^ donor mice into the peritoneal cavity as previously described^28^. In MacGreen mice all monocytes and macrophages in the shed menses-like endometrium are GFP+ ^30^. Two weeks following tissue injection, endometriosis lesions were collected, digested and analysed by flow cytometry. GFP+ macrophages could be detected among Cluster of differentiation (CD)45+, lineage-(CD3, CD19, CD335, Sialic acid-binding immunoglobulin-type lectin F (SIGLEC-F)), Lymphocyte antigen 6 complex, locus G6D (Ly6G)-, Integrin alpha M (CD11b)+ lesion cells (Fig.1A). These data indicate that macrophages derived from the donor endometrium reside within lesions and verifies our previous findings using immunodetection^28^. Endometrial-derived macrophages (GFP+) represented 16% (standard deviation (SD) ± 8.58) of lesion-resident macrophages, whilst the remaining 84% (GFP-; SD ± 8.43) were host derived infiltrating populations (Fig.1B). Next, we sought to determine the origin of the host-derived infiltrating cells. We investigated infiltration of LpM into endometriosis lesions using dual immunodetection for F4/80 (red) and the transcription factor GATA binding protein 6 (GATA6; LpM marker, green; Fig.1C). Quantification of dual positive cells revealed that less than 1% of lesion-resident cells appeared to be derived from LpM (Fig.1D). To verify these findings, we performed adoptive transfer of GFP+ LpM isolated by fluorescent activated cell sorting (FACS) from MacGreen mice (LpM and SpM were determined based on expression of F4/80 and MHCII; see below). GFP+ macrophages could be easily detected in lesions (Fig.1E) suggesting that LpM infiltrate endometriosis lesions and lose expression of GATA6, consistent with a change in phenotype within the lesion microenvironment. Conversely, very few GFP+ macrophages were detected in lesions following adoptive transfer of SpM isolated from MacGreen mice, suggesting that significant trafficking of peritoneal macrophages to lesions was restricted to the LpM population (Fig.1F). Interestingly, GFP+ SpM were instead observed located to the peritoneum adjacent to lesions. Quantification of GFP immunofluorescence revealed that a mean of 27.4% of cells in lesions were derived from LpM and a mean of 3.6% were derived from SpM (Fig1G). To assess infiltration of monocytes into lesions we used dual immunodetection for F4/80 (red) and Lymphocyte antigen 6 complex, locus C (Ly6C; green). We identified Ly6C positive lesion-resident monocytes and a population of dual positive cells (yellow) were also detected indicating that monocytes infiltrated lesions and differentiated *in situ* into macrophages (Fig.1H).

**Figure 1:**
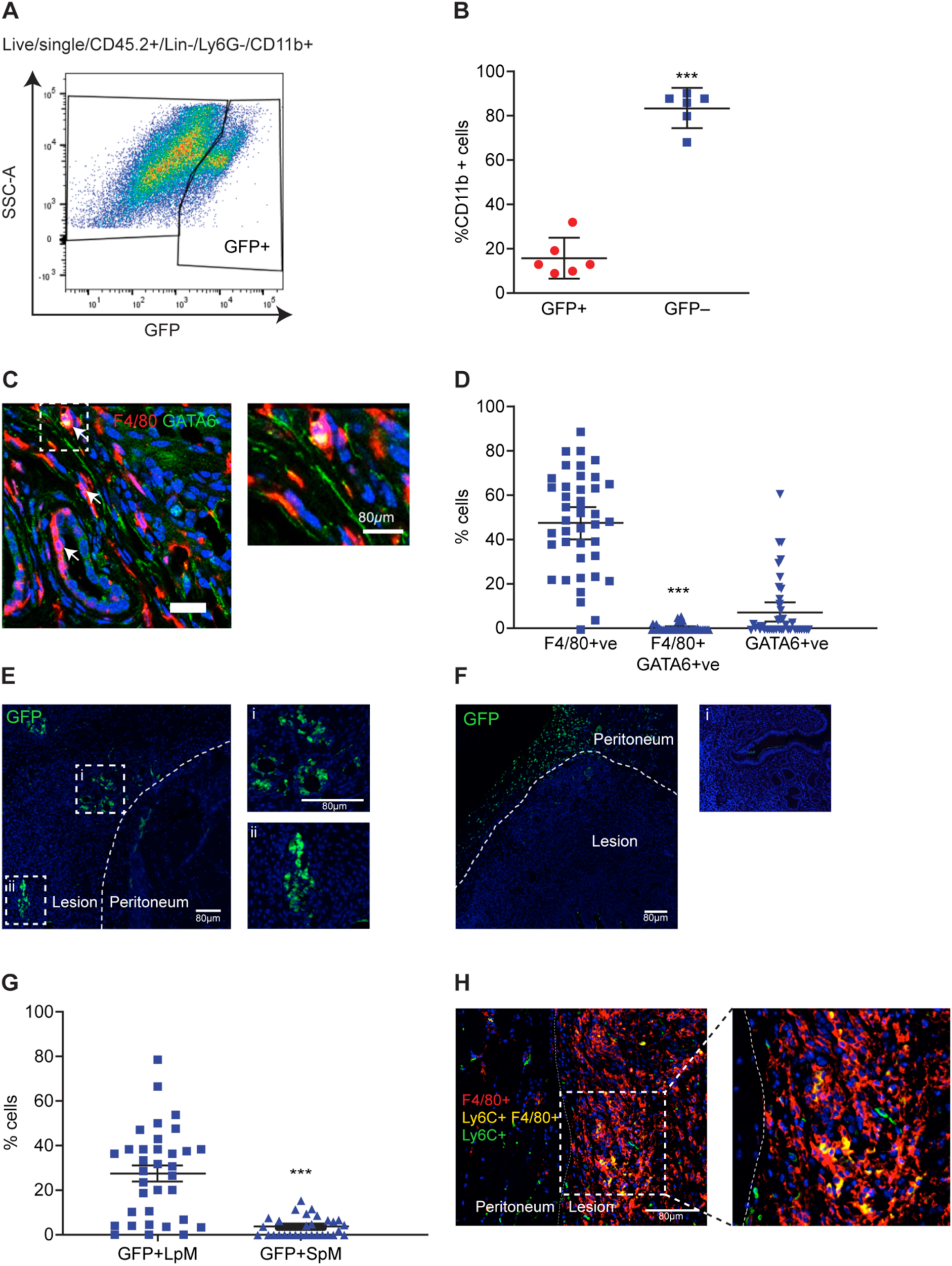
Lesion-resident macrophages have different origins. A) Expression of GFP by lesion-resident macrophages recovered from MacGreen (donor) to wild-type (recipient) endometrial transfers (n=6). B) Quantification of donor endometrial-derived (GFP+) macrophages vs recipient derived (GFP-) macrophages. C) Dual immunodetection for F4/80 (red) and GATA6 (green; n=7 mice (10 lesions)). Thick arrows indicate dual positive cells and thin arrows indicate GATA6-macrophages. D) Quantification of F4/80+, dual positive and GATA6+ cells in lesions. Less than 1% of cells were dual positive for F4/80 and GATA6. E-F) Immunofluorescence for GFP on lesions collected following adoptive transfer of approx. 1×10^6^ LpM (E) or SpM (F) isolated from MacGreen mice. Curved dotted line indicates the boundary between peritoneal and lesion tissue. In E (i) and (ii) show magnified images, in F (i) shows a negative control. G) Quantification of GFP+ LpM and SpM in lesions. H) Dual immunodetection for F4/80 (red) and Ly6C (green) performed on mouse lesions. Data are presented as mean with 95% confidence intervals. Statistical significance was determined using a Student’s t-test. ***;p<0.001.

### Continuous recruitment and contribution of monocytes to peritoneal macrophage populations in mice with induced endometriosis

Next, we investigated macrophage populations present in the peritoneal cavity of mice with induced endometriosis. From CD45+, Ly6G-, lineage-cells, LpM and SpM were determined based on expression of F4/80 and MHCII (Fig.2A). At one-week post tissue injection LpM (F4/80^hi^, MHCII^lo^) numbers were significantly higher (48399 vs 18967 cells / μl, SEM ± 7361) in sham mice (ovariectomised and supplemented with estradiol valerate but given intra-peritoneal (i.p) injection of PBS rather than endometrial tissue; Fig.2B; p<0.01) compared to naïve animals. At 3 weeks, LpM numbers in sham animals had decreased compared to week 1 (p<0.05). In contrast, no significant differences were found in the LpM population in mice with endometriosis. However, a trend can be observed indicating a moderate decrease in LpM numbers at 1 week post-tissue injection in endometriosis mice compared to sham, with an increase at 3 weeks compared to sham. This suggests that there may be some loss of LpM following transfer of endometrial tissue, and this could be attributed to LpM trafficking into lesions or possible clotting of LpM as seen in peritonitis models^31^. By 3 weeks this effect is lost and is consistent with our previous reports demonstrating an increase in LpM at 3 weeks in mice with endometriosis^26^. SpM (F4/80^lo^, MHCII^hi^) numbers were consistent between all groups of animals and time points. Ovariectomy alone can have striking impacts on macrophage pools present in the peritoneal cavity and leads to increased macrophage replenishment^10^. We repeated our results using intact recipient mice and confirmed that in this minimally invasive model, there was no effect on numbers of LpM or SpM suggesting that transfer of endometrial tissue does not disrupt the normal balance of peritoneal macrophages (Fig.S1A). Monocytes in the peritoneal cavity were identified by expression of Ly6C (detection of classical monocytes; Fig.2C) and numbers were significantly increased in mice with induced endometriosis at 1-week post tissue injection compared to naïve mice (Fig.2D; p<0.01). Monocyte numbers remained elevated between weeks 1 and 3 in mice with induced endometriosis, suggesting continuing recruitment to the peritoneal cavity as a consequence of the presence of endometriotic lesions. To begin to investigate the fate of monocytes recruited to the peritoneal cavity of mice with induced endometriosis, we evaluated expression of C-C chemokine receptor type 2 (CCR2; mediates monocyte chemotaxis/ recruitment of monocytes) by F4/80^hi^ (LpM) macrophages in the peritoneal cavity using flow cytometry. Significantly elevated numbers of F4/80+, CCR2+ macrophages were recorded in mice with induced endometriosis (Fig.2E-F), suggesting that increased numbers of monocyte-derived LpM are evident in mice with endometriosis. In steady state conditions, long-lived embryo-derived peritoneal macrophages express T-cell immunoglobulin and mucin domain containing 4 (TIM4), whilst recently monocyte-derived LpM do not^9^, thus to further validate our hypothesis we ascertained TIM4 expression in F4/80^hi^ LpM in intact mice. Contrary to our expectation there was no difference in the ratio of TIM4+ and TIM4-LpM as analysed by flow cytometry at 1-, 2- or 3-weeks post tissue injection (Fig.S1B). We did observe a significant reduction in the proportion of F4/80^lo^, MHCII-, within CD11b+ peritoneal cells between week 1 and week 3-post endometrial tissue injection (Fig.S1C-D). Only a minor proportion of these cells express Ly6C (less than 2% of CD11b cells), indicating that this population may be a transitory state between monocyte and LpM.

**Figure 2:**
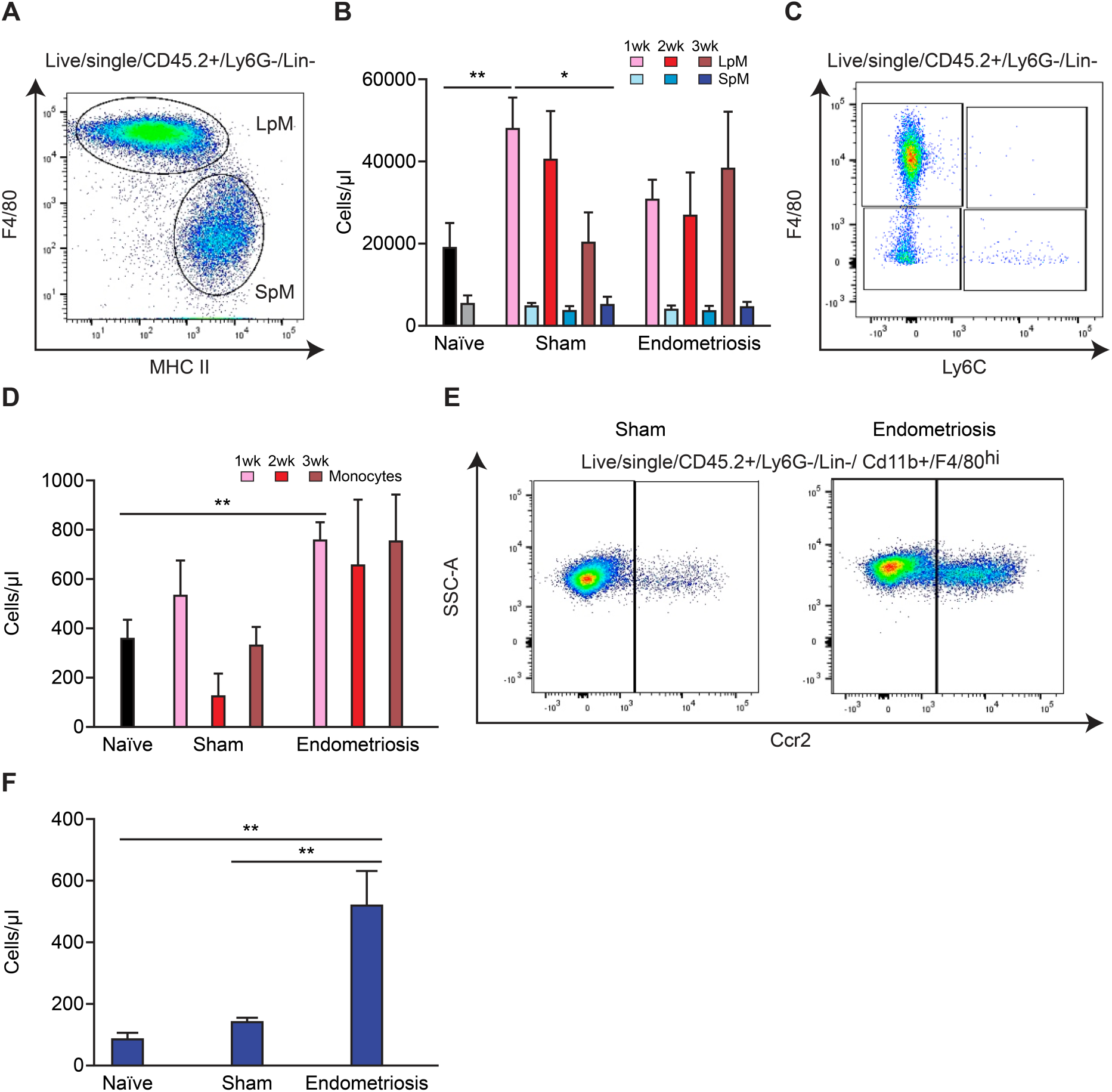
Monocyte recruitment and replenishment of peritoneal macrophage pools in mice with induced endometriosis. A) LpM (F4/80^hi^, MHCII^lo^) and SpM (F4/80^lo^, MHCII^hi^) populations in the peritoneal fluid of mice. B) Quantification of LpM and SpM of sham (n=6-8 each timepoint) and endometriosis mice at 1 (n=6), 2 (n=8) and 3 weeks (n=16) post endometrial tissue injection compared to naïve mice (n=12). C) Flow plot indicating gating of monocytes (F4/80^lo^, Ly6C^hi^) in peritoneal lavage fluid. D) Quantification of monocyte numbers of sham and endometriosis mice at 1, 2, and 3 weeks post tissue injection. E) Flow plot demonstrating expression of CCR2 on F4/80^hi^ macrophages in peritoneal lavage fluid from sham vs endometriosis mice. F) Quantification of CCR2 expression on F4/80^hi^ from endometriosis mice (n=5) compared to sham (n=4) and naïve (n=4) mice. Data are presented as mean ± SEM. Statistics where determined using a one-way ANOVA and a Tukey post-hoc test, *;p<0.05, **;p<0.01.

### Endometrial macrophage depletion leads to reduced endometriotic lesion size

To dissect the role of macrophages with different ontogenies in endometriosis we used a number of depletion strategies. To deplete endometrial macrophages, doxycycline was administered to iCsf1r-KO donor mice from day 15-19 post ovariectomy (Fig.3A), such that we could achieve macrophage depletion by inducibly ablating expression of *Csf1r*^*32*^. Pre-transfer analysis revealed that the endometrium was significantly depleted of F4/80^hi^ macrophages (Fig.3B; p<0.05). Following transfer of macrophage-depleted donor endometrial tissue to wild-type recipients, there was no difference in the number of lesions recovered after two weeks (Fig.3C); however the lesions recovered were significantly smaller than those from mice receiving wild-type endometrium (Fig.3D; p<0.05) with some animals failing to develop any detectable lesions (Fig.3C). These data suggest that macrophages within endometrial tissue promote the growth of endometriosis lesions but do not significantly impact the establishment of lesions.

**Figure 3:**
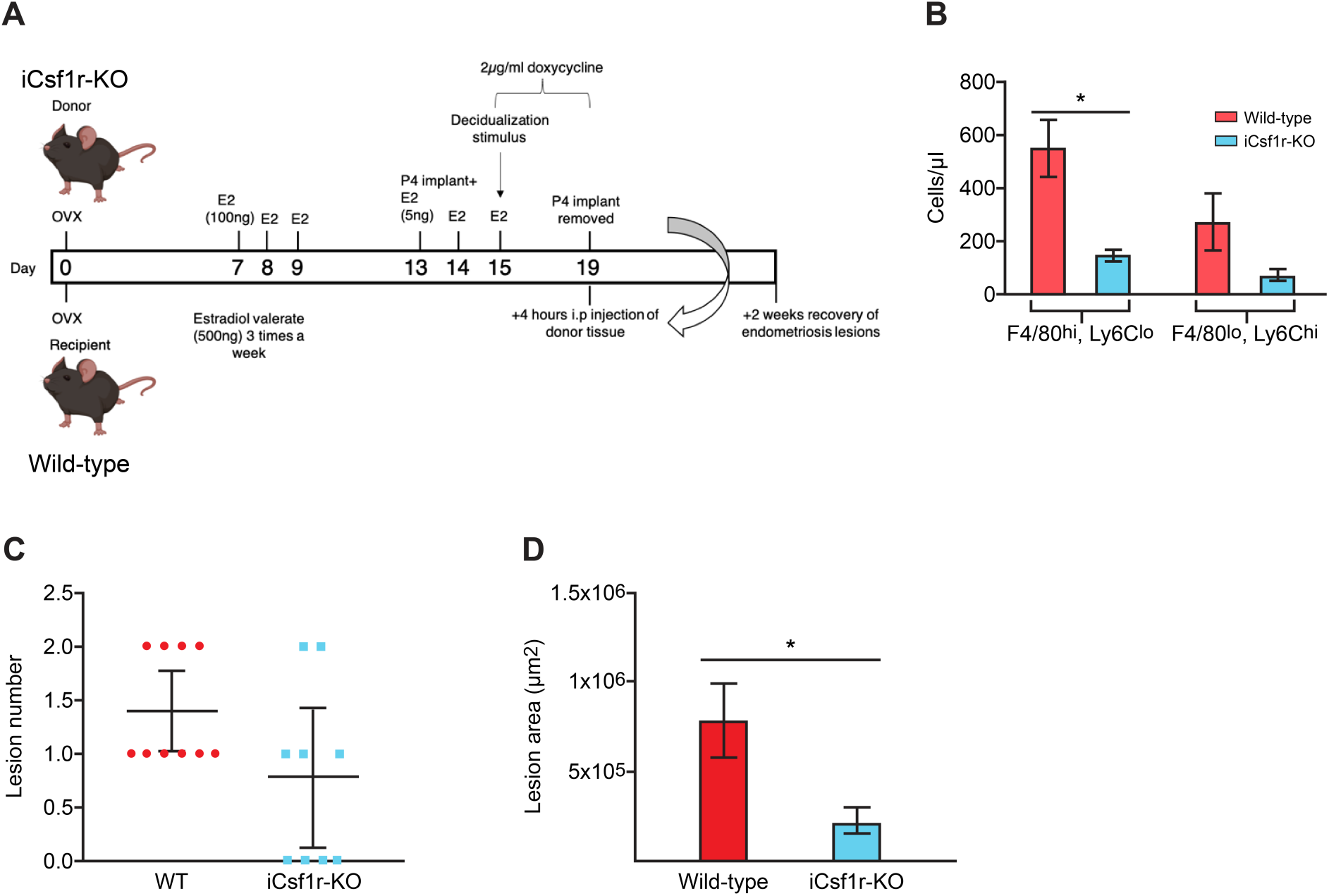
Endometrial macrophage depletion impacts lesion size. A) Schematic demonstrating timing of doxycycline administration to i*Csf1r*-KO donor mice. B) Quantification of F4/80^hi^, Ly6C^lo^ macrophages and Ly6C^hi^, F4/80^lo^ monocytes in donor endometrium from wild-type and iCsf1r-KO mice. C) Number of lesions recovered from wild-type recipient mice receiving either wild-type (n=10) or i*Csf1r*-KO endometrium (n=9). D) Area of lesions recovered from mice receiving either wild-type or i*Csf1r*-KO endometrium. Data are presented as mean ± SEM or 95% confidence intervals (C). Statistical significance was determined using a Student’s t-test. *;p<0.05.

### ‘Monocytopenic’ mice with induced endometriosis had an increased number of lesions

Previous studies have inferred a role for recipient peritoneal macrophages in lesion development since continual ip delivery of clodronate liposomes or anti-F4/80 antibody throughout the growth phase resulted in smaller lesions. Further, ip transfer of bone marrow (BM)-derived macrophages could enhance or inhibit growth dependent on polarisation of macrophages *in vitro* prior to transfer^24^. However, in addition to embryo-derived resident peritoneal macrophages, it is possible these methods also deplete endometrial macrophages (in transferred endometrial tissue) and recruited monocyte-derived cells. Notably, these studies showed that liposome-mediated depletion of embryonic resident peritoneal macrophages prior to endometrial transplantation did not significantly effect lesion development^24^, suggesting the role of embryonic LpM is, at best redundant. Hence, we next aimed to prevent recruitment of host monocytes to the peritoneal cavity and lesions using *Ccr2* null monocytopenic recipient mice. LpM, SpM and monocytes were all significantly reduced in frequency (4.1×10^5^ vs 7.9×10^5^, 1.1×10^5^ vs 2.8×10^5^ and 1.3×10^4^ vs 3.6×10^4^ respectively) in the peritoneal lavage fluid of *Ccr2*-/- mice with induced endometriosis (Fig.4A-E and FigS2A-C). This supports the concept that in our mouse model of endometriosis monocytes are continually recruited and contribute to the LpM and SpM pools. Furthermore, while lesion size was not different between the two groups (Fig.4G), *Ccr2*-/- mice had significantly more lesions (p<0.05) compared to wild-type mice (Fig.4F). This indicates that monocytes or monocyte-derived macrophages normally protect against establishment of lesions. To validate these findings, we repeated the experiment in Chemokine (C-C motif) ligand 2 null (*Ccl*2-/-) mice. In *Ccl2*-/- mice with endometriosis, LpM and SpM were depleted (6.8×10^4^ vs 1.4×10^5^, 8.4×10^3^ vs 3.7×10^4^ respectively) in the peritoneal lavage compared to wild-type mice (Fig.5A-C; p<0.05 and Fig.S2D-F). We observed a concomitant increase in the number of lesions recovered from *Ccl2*-/- mice (Fig.5E; p<0.05). Strikingly, *Ccr2*-/-and *Ccl2*-/- mice with induced endometriosis were still able to recruit Ly6C+ monocytes from the bone marrow to the peritoneal cavity despite monocytes being absent in the peritoneal lavage of naïve and sham monocytopenic mice (Fig.S3). Moreover, in lesions recovered from *Ccr2*-/- and *Ccl2*-/- mice we could still detect monocytes using immunodetection (Fig.4H-I and Fig.5G-H) indicating that there is some redundancy in the CCL2-CCR2 axis in the presence of endometriosis lesions and suggests that monocytes may be recruited directly from the bone marrow via another mechanism.

**Figure 4:**
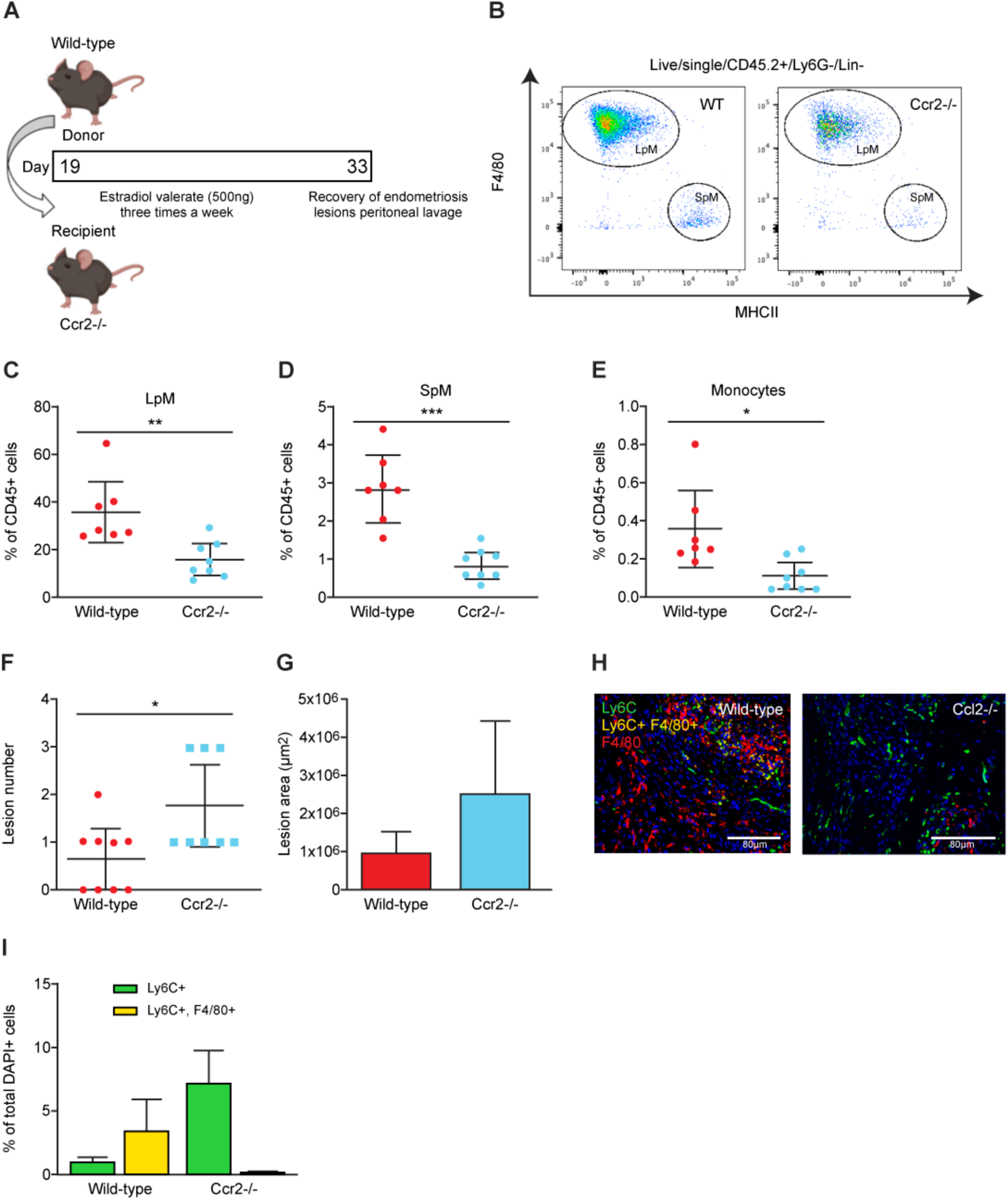
‘Monocytopenic’ mice with induced endometriosis establish more lesions. A) Schematic demonstrating experimental design. Wild-type donor endometrium was generated as previously shown (Fig.3A) and injected ip into ovariectomised *Ccr2*-/- recipients. Wild-type recipients were also used as controls. B) Flow plot indicating gating and number of LpM (F4/80^hi^) and SpM (MHCII^hi^) in peritoneal lavage fluid recovered from wild-type (n=7) and *Ccr2*-/- mice (n=8) with induced endometriosis. C-E) Quantification of (C) LpM, (D) SpM, and (E) monocytes (Ly6C^hi^) in peritoneal lavage fluid. F) Number of lesions recovered from wild-type and *Ccr2*-/- mice with induced endometriosis. G) Size of lesions recovered from wild-type and *Ccr2*-/- mice with induced endometriosis. H) Dual immunodetection for Ly6C (green) and F4/80 (red) on lesions recovered from wild-type and Ccr2-/- mice I) Quantification of monocytes (Ly6C+; yellow bars) and monocyte-derived macrophages (Ly6C+, F4/80+; green bars) in lesions recovered from wild-type and *Ccr2*-/- mice. Data are presented as mean ± SEM or 95% confidence intervals (F). Statistical significance was determined using a Student’s t-test or Mann-Whitney test. *;p<0.05, **;p<0.01, **;p<0.001.

**Figure 5:**
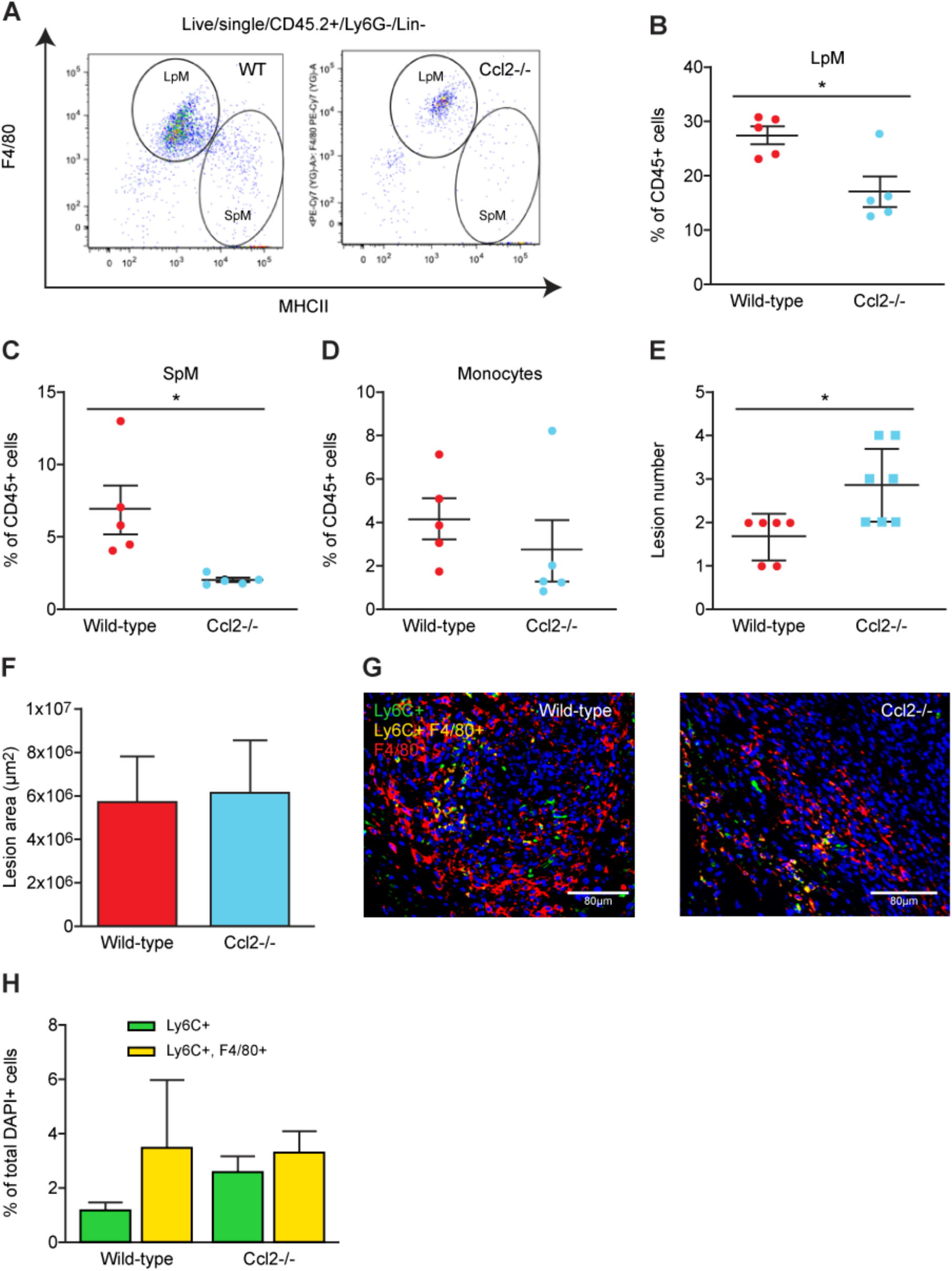
More lesions are evident in *Ccl2*-/- mice. A) LpM and SpM populations in peritoneal lavage fluid from wild-type and *Ccl2*-/- mice with induced endometriosis. (B-D) Quantification of (B) LpM,(C) SpM, and (D) monocytes in peritoneal lavage fluid from wild-type (n=5) and *Ccl2*-/- (n=5) mice with induced endometriosis. E) Number of lesions recovered from wild-type and *Ccl2*-/- mice. F) Size of lesions recovered from wild-type and *Ccl2*/- mice with induced endometriosis. G) Dual immunodetection for Ly6C (green) and F4/80 (red) on lesions recovered from wild-type and *Ccl2*-/- mice H) Quantification of monocytes (Ly6C+) and monocyte-derived macrophages (Ly6C+, F4/80+) in lesions recovered from wild-type and *Ccl2*-/- mice. Data are presented as mean ± SEM. Statistical significance was determined using a Student’s t-test or Mann-Whitney test. *;p<0.05.

### Transient depletion of monocytes did not impact establishment of endometriotic lesions

Next, we sought to transiently deplete monocytes to ascertain the role of undifferentiated monocytes in lesion development whilst leaving the peritoneal macrophage populations relatively unaltered. We depleted monocytes using a function blocking CCR2 monoclonal antibody (MC21) injected into the peritoneal cavity of mice with induced endometriosis. Mice were injected with MC21, 6 hours prior to transfer of endometrial tissue, and every following day for 4 days (Fig.6A). Compared to a control IgG antibody (MC67), MC21 triggered an expansion of LpM (Fig.6C; p<0.05), did not modify SpM (Fig.6D) and significantly reduced the number of monocytes in the peritoneal cavity (Fig.6B and E; p<0.05). In mice with induced endometriosis treated with MC21 there was no difference in lesion number or size compared to mice treated with control MC67 (Fig.6F-G). These results rule out the possible contribution of recently recruited monocytes to the ‘anti-endometriosis’ function of monocyte-derived cells observed in Fig.4-5.

**Figure 6:**
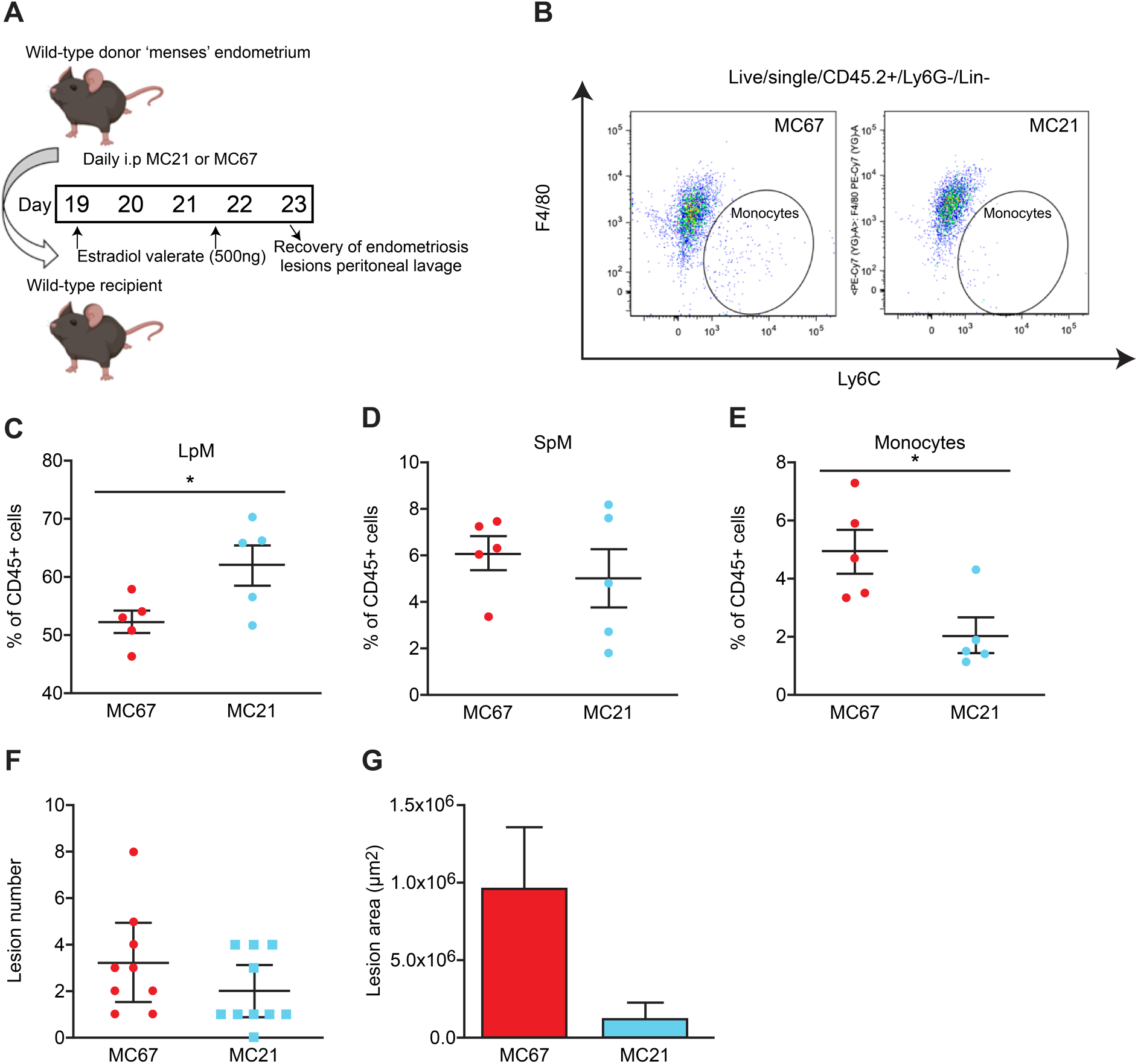
A function-blocking Ccr2 mAb reduces monocyte numbers without significant impact on lesion number. A) Schematic showing the experimental design; mice with induced endometriosis were treated with a control IgG (MC67; n=9) or a function blocking CCR2 mAb (MC21; n=10). B) Flow plot demonstrating the numbers of Ly6C^hi^ monocytes in mice with induced endometriosis treated with MC67 or MC21. C-E) Quantification of (C) LpM, (D) SpM, (E) monocytes in the peritoneal lavage fluid of mice with induced endometriosis. F) Number of lesions recovered from mice treated with MC67 or MC21. G) Size of lesions recovered from mice treated with MC67 or MC21. Data are presented as mean ± SEM. Statistical significance was determined using a student’s t-test or a Mann-Whitney test. *;p<0.05.

## Discussion

The pathophysiology of endometriosis remains enigmatic^21^. Although immune cell dysfunction is intrinsically linked with the disorder, our understanding of macrophage origins and respective function remains limited compared with other diseases, such as cancer^34^. Macrophages are critical for endometriotic lesion establishment and survival, as indicated by clodronate liposome and anti-F4/80 antibody mediated depletion experiments^24^. They play a key role in promoting growth and vascularization of ectopic tissue^25^ and in encouraging nerve growth, activation of nerves and pain generation in endometriosis^26, 27^. As the ontogeny of macrophages in diseased tissue is a key determinant of how they respond and contribute to pathogenesis, it is necessary to understand how macrophages derived from different sources impact lesion development. This will provide a greater appreciation of how ‘disease-promoting’ or ‘protective’ macrophages could be targeted in the future as a potential therapy.

The use of a syngeneic mouse model of induced endometriosis^28^ has allowed intricate exploration of macrophage ontogeny and functional consequences of depleting specific populations that is impossible to perform in women. In this study, we have used a model that aims to recapitulate the process of ‘retrograde menstruation’ by generating ‘menses’-like endometrium in donor mice and injecting the tissue into the peritoneal cavity of recipient mice. Both lesion number and size were recorded as a determinant of macrophage role in either establishment or growth of lesions, respectively. We have previously demonstrated that lesion-resident macrophages are derived from both the donor endometrium and the recipient^28^. In the current study we confirm this observation and further establish that host-derived macrophages originate from infiltrating LpMs and from monocytes. We used different depletion strategies to explore the impact of macrophages from different origins on lesion development.

The contribution of embryonic-derived and monocyte-derived macrophages to the ‘tissue-resident’ population is different in each tissue^3, 4, 5^. We previously demonstrated that macrophages present in the endometrium can be detected in lesions recovered from our model of induced endometriosis^28^. However, little is known regarding the ontogeny of macrophages in the endometrium. In a mouse model of menstruation, utilizing MacGreen (*Csf1r-EGFP*) mice, Cousins et al demonstrated that three populations could be distinguished in the endometrium using dual staining for F4/80 and GFP: (1) a population of GFP+, F4/80-cells likely to be infiltrating monocytes, (2) a population of GFP+, F4/80+ cells suggestive of monocyte-derived macrophages, and finally (3) a population of putative (but not confirmed with lineage tracing) ‘tissue-resident’ macrophages that are GFP-, F4/80+. The cells were localized to areas of breakdown, repair and remodelling, respectively^30^. The ‘menses-like’ endometrium that we recover from donor mice for transfer into recipient mice is collected at the initiation of the ‘break-down’ phase and is most likely to consist of monocytes and monocyte-derived macrophages. Moreover, the tissue collected is the ‘decidual’ mass only and does not include the compartment of the uterus where putative ‘tissue-resident’ macrophages are located. Using an inducible *Csf1r*-knockout to generate donor endometrium we achieved depletion of macrophages (F4/80^hi^, Ly6C^lo^) and limited the number of monocytes (F4/80^lo^, Ly6C^hi^) in the tissue transferred to recipient mice. We did not observe any difference in the number of lesions formed between recipient mice that received wild type or macrophage depleted endometrium, however we did find that the lesions recovered were significantly smaller in mice receiving macrophage-depleted endometrium, indicating that endometrial macrophages play a critical role in growth of lesions. The breakdown phase of the (donor) endometrium is analogous to the initial inflammatory phase of the wound healing process where pro-inflammatory macrophages play a vital role prior to wound repair^35^. If we consider endometriotic lesions as chronic wounds that do not fully resolve their inflammation, we may presume that the initial inflammatory phase begins as the endometrium breaks down during menstruation (or within the established lesion during cyclical remodelling) and the repair of the translocated endometrium occurs in the peritoneal cavity resulting in the formation of lesions. The subsequent phase of the tissue repair process is the proliferative phase: in mouse models of skin injury, macrophage depletion during this phase resulted in granulation tissue that had very few blood vessels and proliferative cells and a significant reduction in wound closure. This indicates that macrophages support this phase by promoting endothelial cell survival and vascularization which facilitates proliferation^36^. We suggest that a similar process occurs in mice receiving endometrium-depleted of monocytes and macrophages and this may explain reduced lesion size in recipient mice.

The ontogeny of peritoneal macrophages is better characterized. LpM are embryo-derived, long-lived and undergo self-renewal, however monocytes do continually enter the peritoneal cavity via CCR2 where they continually replenish the SpM compartment and infrequently differentiate into LpM^9^. Such replenishment of LpM occurs in a sexually dimorphic manner, occurring more quickly in male compared to female mice who retain their embryo-derived LpM for much longer^9, 15^, and it is likely that in adult female mice of the age used in our studies (10-12 wks) only 10-30% of peritoneal LpM would be derived from adult monocytes^10, 15^. In this study we used ovariectomy and estradiol supplementation of recipient animals to allow optimal lesion development. Peritoneal surgery increases the contribution of monocytes to the peritoneal LpM however; the embryonic component would still be expected to comprise approximately 50% of the population at the time of injection of endometrial material. Although monocyte-derived LpM mostly phenocopy embryo-derived LpM, transcriptomically they exhibit some differences. For example, embryo-derived LpM express *Timd4*, whilst in steady state many monocyte-derived LpM do not, or take significant time to do so^9, 10^. Notably, the number of monocytes and CCR2+ LpM was significantly elevated in the peritoneal cavity of mice with induced endometriosis, above levels seen in ovariectomised controls, suggesting monocytes are continually recruited and contribute to LpM pools during lesion development.

Consistent with a significant input of monocytes to LpM in endometriosis, our most striking result was revealed by constitutively limiting monocyte recruitment (using both *Ccr2*-/- and *Ccl2*-/- mice) and subsequently reducing both LpM and SpM replenishment, which resulted in mice with induced endometriosis developing significantly more lesions. Thus, we suggest that monocyte-derived macrophages act to protect the peritoneal cavity and can limit the establishment of lesions (Fig.7). When left unmanipulated, female *Ccr2*-/- mice normally exhibit equivalent numbers of LpM to wild-type controls^9^. Despite this input, the frequency of TIM4+ and TIM4-LpM did not change between weeks 1- and 3-post tissue injection, suggesting that a mechanism exists to drive rapid expression of TIM4 on monocyte-derived LpM during endometriosis. These may be similar to those that underlie the rapid acquisition of TIM4 by monocyte-derived LpM recently described to occur in the pleural cavity during helminth infection^37^. Thus, in endometriosis we have demonstrated heightened monocyte input into LpM and potential pathology-associated acquisition of TIM4 by monocyte-derived LpM.

**Figure 7:**
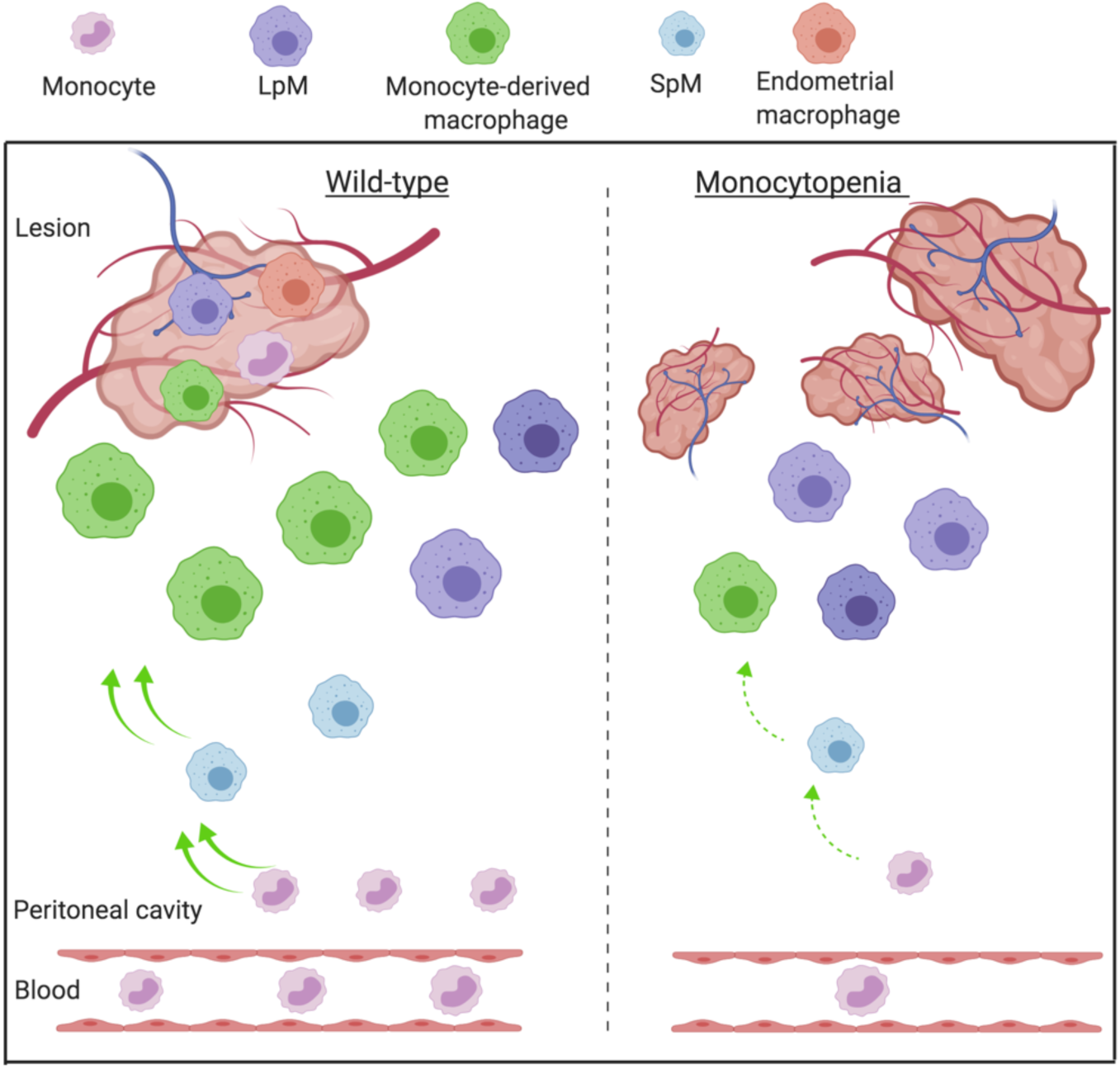
Monocyte-derived macrophages are guardians of the peritoneal cavity in mice with induced endometriosis. Lesion-resident macrophages are a heterogenous population constituted by macrophages that have different origins; endometrium, peritoneum, and recruited monocytes that differentiate into macrophages in lesions. Wild-type mice with induced endometriosis have increased monocyte recruitment and replenishment of SpM and LpM pools by monocyte-derived cells. In mice where monocyte recruitment is constitutively limited (*Ccr2*-/- or *Ccl2*- /-), LpM and SpM pools are significantly reduced suggesting that the majority of LpM in the peritoneal cavity are embryo-derived. Elevated numbers of lesions develop in mice with constitutively reduced monocyte numbers suggesting that monocyte-derived macrophages protect the peritoneal cavity when challenged with ectopic endometrial tissue. We propose a putative model where endometrial macrophages promote lesion growth, whilst monocyte-derived macrophages (possibly monocyte-derived LpM) protect the peritoneal cavity against establishment of lesions.

In monocytopenic mice with endometriosis, a reduction in SpM, LpM as well as monocytes was observed. Monocytopenic mice also exhibited significantly more lesions, indicating that monocyte-derived cells are ‘anti-endometriosis’, although to which population of monocyte-derived cells this function can be attributed remains uncertain. As transient antibody-mediated depletion of monocytes during the establishment phase failed to increase the number of lesions but also left SpM and LpM populations intact, it seems likely that either SpM or LpM provide a dominant anti-endometriosis affect.

We determined that a significant proportion of LpM (27.4% of lesion resident cells were GFP+ following adoptive transfer of LpM) but not SpM enter endometriosis lesions using adoptive transfers; we suggest that these LpM change phenotype in lesions very rapidly because only a few lesion cells positive for both F4/80 and GATA6 (<1%) were identified using dual staining, whereas LpM in the cavity are almost entirely positive for GATA6^11^. Aside from identifying peritoneal LpM as a source of macrophages in endometriosis lesions, these findings are important because they show that LpM are able to re-programme and survive in ectopic tissue, a topic of significant controversy in the field of tissue-resident macrophage biology^38, 39^. Indeed, our results are consistent with the reversible expression of GATA6 by LpM in the absence of sustained retinoic acid receptor signalling^11^ and mirror recent findings that pericardial cavity GATA6+ macrophages lose expression of GATA6 following recruitment to areas of ischemic heart disease^40^. In the same manner, mature F4/80^hi^ GATA6+ peritoneal LpM are reported to traffic directly across the mesothelium into the liver following sterile injury. Once in the liver, the macrophages undergo local proliferation, and up regulation of markers of alternative activation, such as Arginase 1 (Arg1) and Retnla. In the absence of peritoneal macrophages, healing was significantly delayed^41^. Our data suggest that in endometriosis LpM trafficking to lesions may play a similar role, perceiving the ectopic tissue as a wound and activating repair processes. Interestingly, GATA6 positive macrophages that invade the epicardium from the pericardial space following experimental myocardial infarction were anti-fibrotic, despite a rapid loss of GATA6 expression^40^. Fibrosis is a consistent feature of endometriotic lesions^42^. Whilst eutopic endometrium is able to undergo scar-free healing to restore full tissue functionality, when the tissue is translocated to the peritoneal environment fibrotic ‘repair’ occurs to form lesions. The mechanisms responsible for this are yet to be fully resolved but pro-repair macrophages have been implicated in the process^43^. Thus, it seems unlikely that LpMs trafficking into lesions contribute to fibrotic repair, however, the ontogeny of macrophages contributing to fibrogenesis in endometriosis remains to be determined. Interestingly, our data suggest that SpM (which likely include both classical dendritic cells (cDC)1 and cDC2, with the flow cytometry gates used in our study) may have a neutral role in the pathophysiology of endometriosis since they neither increased in number nor appeared to significantly contribute to the lesion-resident population. Hence, we suggest it is likely that the monocyte-derived LpM are protective against development of endometriosis.

Our data indicates a key role for monocyte recruitment to the peritoneal cavity *and* ectopic tissue in endometriosis. We readily detected lesion-resident monocytes as well as cells that were double positive for both Ly6C and F4/80, but whether this indicates that monocytes rapidly differentiate in lesions or that differentiated monocyte-derived macrophages are recruited from the peritoneal cavity remains unclear. We speculate that monocyte-derived cells represent the largest lesion-resident population. However, we cannot currently conclude whether these are monocyte-derived LpM or monocytes recruited directly to the lesion. If the latter, we do not know whether these come via the cavity or through the newly formed vasculature associated with the lesion.

In a previous study, Bacci et al depleted peritoneal macrophages using liposomal clodronate in a mouse model of endometriosis. Depletion of peritoneal macrophages prior to transfer of endometrial tissue had no significant impact on lesion establishment and growth^24^ implying that embryonic-derived LpM are redundant in lesion development. Continuous depletion during establishment and growth resulted in significantly smaller lesions, allowing the authors to conclude that the dominant role for macrophages in endometriosis is to promote the development of lesions. It may be presumed that this approach has the potential to deplete all monocytes and macrophages including those in the transferred endometrial tissue. Thus, in light of our findings that donor endometrial macrophages play a significant role in facilitating lesion growth we suggest the findings of Bacci et al are a consequence of global macrophage depletion. In contrast, our study used depletion strategies that target macrophages from different origins. We demonstrate a protective role for monocyte-derived macrophages (presumably in LpM form). Similarly, further experiments by Bacci et al demonstrated that ip transfer of bone marrow (BM)-derived macrophages could enhance or inhibit growth dependent on polarisation of macrophages *in vitro* prior to transfer^24^. Further studies will aim to elucidate the exact phenotype and mechanism of monocyte-derived LpM in endometriosis.

Definition of the macrophage populations that reside in diseased tissues is vital for understanding macrophage-driven pathology, particularly for endometriosis where little is known about the lesion macrophage niche. In the future it may be possible to harness the protective properties of monocyte-derived macrophages as a potential therapy for women with endometriosis. We propose a putative model that in endometriosis, macrophages derived from the endometrium exhibit ‘pro-endometriosis’ functions and facilitate growth of endometriotic lesions, whereas monocyte-derived cells, possibly in the form of LpM from the cavity have an ‘anti-endometriosis’ role and are protective against persistence of ectopic tissue and establishment of lesions.

Collectively, we have demonstrated multiple origins for endometriotic lesion-resident macrophages, a key role for monocyte-derived macrophages in protecting the peritoneal cavity when challenged with ectopic endometrial tissue and a pathological role for endogenous endometrial macrophages. Our findings imply that monocyte recruitment or monocyte-derived macrophages may be defective in women with endometriosis, and this should be explored in more depth in women with the condition. Thus, we have opened up new avenues and possibilities for how macrophages can be targeted or harnessed as a therapeutic option in the treatment of endometriosis.

## Materials and Methods

### Animals and reagents

Wild-type C57BL/6JOIaHsd female mice were purchased from Harlan (Harlan Sprague Dawley Inc, Bicester, UK) at 8-12 weeks of age. All transgenic lines used in this study were on the C57BL/6 background, and were bred and maintained at the University of Edinburgh or the University of Warwick. All animal work was licensed and carried out in accordance with the UK Home Office Animal Experimentation (Scientific Procedures) Act 1986 and the work licensed under PPL 70/8731 (E.G). Mice had access to food and water ad libitum and were kept at an ambient temperature and humidity of 21°C and 50% respectively. Light was provided 12 hours a day from 7am-7pm. *ROSA26-rtTA:tetO-Cre:Csf1rflox/flox*^32^ (colony stimulating factor 1 receptor (Csf1r) conditional knock out) allows deletion of *Csf1r* following treatment with the tetracycline analog doxycyclinehyclate (2μg/ml in 5% sucrose water; Merck) causing CSF1R expressing macrophage populations to be depleted. *B6*.*Cg-Tg(Csf1r-EGFP)1Hume/J* (MacGreen) express enhanced green fluorescent protein (EGFP) under control of the *Csf-1r* promoter^44^. We used a number of strategies to selectively deplete different monocyte-derived populations in the peritoneal cavity and in lesions; 1) *B6*.*129S4-Ccr2tm1Ifc/J (C-C chemokine receptor type 2 (Ccr2 -/-)* mice have a homozygous mutation in the *Ccr2* gene^45^. They have reduced monocytes, monocyte-derived macrophages and small peritoneal macrophages in the peritoneal cavity in addition to a low number of circulating Ly6C^hi^ monocytes due to an inability for monocytes to extravasate from the bone marrow and from blood vessels. 2) *B6*.*129S4-Ccl2tm1Rol/J (Chemokine (c-c motif) ligand 2 (Ccl2) -/-)* mice possess a mutation in the *SCYA2* gene encoding the CCL2 ligand. *Ccl2*-/- mice have normal peritoneal macrophage numbers but reduced recruitment of monocytes and monocyte-derived macrophages into the peritoneal cavity under inflammatory conditions^46^. 3) The monocyte depleting rat anti-mouse CCR2 mAb (clone MC21) isotype IgG2b^47^ was used for monocyte depletion experiments. Mice received a daily intraperitoneal injection of 20μg per mouse of MC21 to prevent infiltration of monocytes into the peritoneal cavity. Isotype-matched rat IgG2b control antibody (MC67) was used as a control.

### Mouse model of induced endometriosis

Endometriosis was induced in mice using a syngeneic model as previous described^28^. The model aims to mirror the process of ‘retrograde menstruation’. In brief, donor mice were induced to undergo a ‘menses’-like event by removing the ovaries and exposing the mice to a hormonal schedule similar to a truncated menstrual cycle and a stimulus that causes the endometrial stromal cells to undergo decidualization^48^. Following P4 withdrawal the endometrial lining begins to shed. 4-6hrs after withdrawal of P4 the ‘menses’-like endometrium is collected and injected into ovariectomized mice supplemented with estradiol valerate^48^. Lesions are recovered that contain stoma +/- epithelial cells and immune cell influx^28^. Unless otherwise stated lesions were collected 2 weeks following tissue injection. For clarity, individual experiments using different combinations of transgenic and wild-type mice are described: *Experiment 1: Endometrial macrophage incorporation into lesions*. Endometrium from MacGreen donors was injected i.p into wild-type C57BL/6 recipients (n=6 mice). *Experiment 2: Incorporation of peritoneal macrophages into lesions*. LpM (F4/80^hi^, MHCII^lo^) or SpM (F4/80^lo^, MHCII^hi^) were isolated from MacGreen mice using fluorescent activated cell sorting (FACs) and adoptively transferred into the peritoneal cavity of recipient mice (n=12 per population) at the same time as donor endometrium, both donor and recipient were wild-type C57BL/6. Incorporation was evaluated using immunodetection of GFP. *Experiment 3: Impact of endometrial macrophage depletion on lesion formation*. Macrophages were depleted in donor endometrium by administering doxycycline to *Csf1r*-cKO mice between days 15-19 of the ‘menses’ protocol (Fig.4A). Macrophage depleted endometrium was injected i.p into wild-type recipients (n=9). For comparison wild-type endometrium was transferred into wild-type recipients (n=10). *Experiment 4: Depletion of monocytes in the peritoneal cavity (i)*. Donor endometrium from wild-type C57BL/6 mice was injected i.p into wild-type (control; n=7) or *Ccr2*-/- recipients (n=8). *Experiment 5: Depletion of monocytes in the peritoneal cavity (ii)*. Wild-type donor endometrium was injected i.p into wildtype (controls; n=5) or *Ccl2*-/- recipients (n=5). In some experiments we used recipients that had not been ovariectomized (intact). Endometrial tissue was generated and injected in the same way as the standard model. Recipient mice did not receive any hormonal manipulation. In both models, mice were culled 14 days post tissue injection (unless otherwise stated) and endometriosis lesions and peritoneal lavage were collected. Peritoneal lavage was recovered by injecting 7 ml ice-cold DMEM into the peritoneal cavity followed by gentle massage and recovery. Lesions were either collected into neutral-buffered formalin for paraffin embedding and immunohistochemical analysis or DMEM for flow cytometry analysis.

### Flow cytometry

Endometrium or lesions were dissected, pooled from each mouse and placed in 2ml ice-cold DMEM. Tissues were cut into small pieces using a scalpel and digested with 1 unit of Liberase DL, 1 unit of Liberase TL and 0.6 mg DNAse enzymes. The tissue and enzymes were incubated for 45 minutes at 37°C, with vortexing every 5 minutes. Following digestion samples were filtered through 100μM filters. Red blood cells were lysed from peritoneal lavages and cells derived from the endometrium or lesions, and approx. 10^6^ cells per sample were blocked with 0.025mg anti-CD16/32 (clone 93; BioLegend, San Diego, CA, USA) and then stained with a panel of antibodies shown in Table 1. Brilliant™ violet stain buffer was included when required. Fluorescence minus one (FMO) and unstained controls were used to validate gating strategies. Just prior to analysis on the flow cytometer, DAPI and 123count eBeads (Thermo Fisher Scientific) were added to samples. Samples were analysed using an LSRFortessa with FACSDiva software (BD Biosciences) or FACSMelody with Chorus software and analysed with FlowJo v.9 software (FlowJo, Ashland, OR, USA). Analysis was performed on single, live cells determined using forward scatter height vs. area and negativity for live/ dead (DAPI or alternative viability dye). For fluorescent activated cell sorting, red blood cell lysis, Fc blocking and fluorescent staining was performed as previously described and samples sorted into pure cell populations based on cell surface marker expression using a FACS Fusion (BD Biosciences). For data where peritoneal populations are expressed as cells/ μl, absolute counts were calculated using 123 eBeads and the equation: absolute count (cells/ µl) = (cell count / eBead count) x eBead batch concentration. Final volume for cytofluorimetric analysis was 300 μl. Thus for example, 400 cells / μl is equal to 1.2 x10^5^ cells per cavity

**Table 1.**
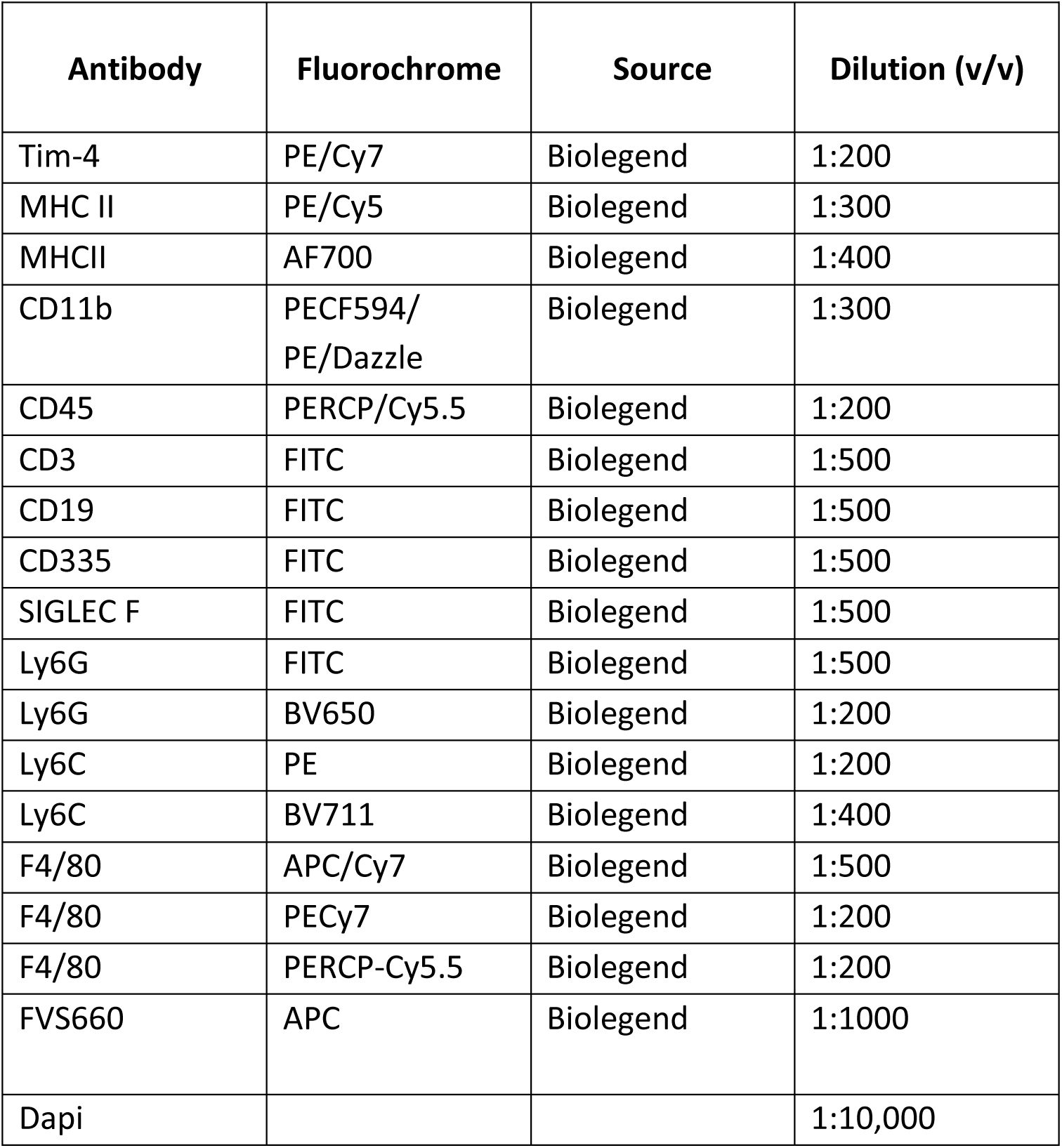
Flow cytometry antibodies.

| Antibody | Fluorochrome | Source | Dilution (v/v) |
| --- | --- | --- | --- |
| Tim-4 | PE/Cy7 | Biolegend | 1:200 |
| MHC II | PE/Cy5 | Biolegend | 1:300 |
| MHCII | AF700 | Biolegend | 1:400 |
| CD11b | PECF594/<br>PE/Dazzle | Biolegend | 1:300 |
| CD45 | PERCP/Cy5.5 | Biolegend | 1:200 |
| CD3 | FITC | Biolegend | 1:500 |
| CD19 | FITC | Biolegend | 1:500 |
| CD335 | FITC | Biolegend | 1:500 |
| SIGLEC F | FITC | Biolegend | 1:500 |
| Ly6G | FITC | Biolegend | 1:500 |
| Ly6G | BV650 | Biolegend | 1:200 |
| Ly6C | PE | Biolegend | 1:200 |
| Ly6C | BV711 | Biolegend | 1:400 |
| F4/80 | APC/Cy7 | Biolegend | 1:500 |
| F4/80 | PECy7 | Biolegend | 1:200 |
| F4/80 | PERCP-Cy5.5 | Biolegend | 1:200 |
| FVS660 | APC | Biolegend | 1:1000 |
| Dapi |  |  | 1:10,000 |

### Immunofluorescence

Immunofluorescence was performed as previously described^26, 27, 49^. In brief, sections were antigen retrieved with heat and pressure (buffers pH 6.0 or pH 9.0) or trypsin tablets dissolved in dH2O (for F4/80 antibody; Sigma) and incubated with sections for 20 min at 37°C. Sections were blocked for endogenous peroxidase (6% H_2_O_2_ in methanol) and nonspecific epitopes (species-specific serum diluted 1:5 in Tris-buffered saline and 5% bovine serum albumin, or blocking serum from species specific ImmPRESS^®^ kit; Vector Laboratories) and incubated with primary antibody (Table 2) at 4°C overnight. Antibody detection was performed using a secondary antibody conjugated to horseradish peroxidase, often from an ImmPRESS^®^ polymer detection kit followed by colour development using a tyramide signal amplification system kit with cyanine (Cy)3 or fluorescein (1:50 dilution; PerkinElmer, Waltham, MA, USA). For detection of the second antigen in dual immunofluorescence, sections were boiled in citrate buffer, and the second primary antibody applied overnight and detected as above. Nuclei were stained with DAPI and sections mounted in Permafluor (Thermo Fisher Scientific). Images were captured on a LSM710 confocal microscope and AxioCam camera (Carl Zeiss). Mouse uterus was used as a positive control tissue, and negative controls had omission of the primary antibody

**Table 2.** Antibodies for immunodetection.

| Antibody | Source | Cat number | Target cell | Species raised | Dilution (v/v) | Secondary used |
| --- | --- | --- | --- | --- | --- | --- |
| F4/80 | eBioscience | 14-4801 | Macrophage | Rat | 1:600 | ImmPRESS®<br>HRP-conjugated<br>anti-rat<br>antibody |
| GATA6 | Cell<br>Signalling<br>Technology | 5851S | LpM | Rabbit | 1:3000 | ImmPRESS®<br>HRP-conjugated<br>anti-rabbit<br>antibody |
| Ly6C | Abcam | Ab15627 | Monocytes | Rat | 1:100 | ImmPRESS®<br>HRP-conjugated<br>anti-rat<br>antibody |
| GFP | Abcam | Ab6556 | Macrophages<br>labelled with<br>EGFP | Mouse | 1:2000 | ImmPRESS®<br>HRP-conjugated<br>anti-mouse<br>antibody |

### Fiji analysis

For cell counting of F4/80, GATA6 dual immunofluorescent stains, 4 random images at x63 objective were taken from each lesion and images quantified using Fiji plugin ‘Cell Counter’. Total nuclei were counted, as well as cells positive for respective markers. Values were expressed as % of total DAPI+ cells. The area of H&E stained lesions captured using 2.5x magnification was measured in Fiji by setting the scale to a known size (198 pixels = 500μM) and then drawing around boundary of the lesion (excluding peritoneal and adipose tissue) and using the ‘measure’ function.

### Definiens analysis

Ly6C, F4/80 dual immunofluorescence was automatically quantified using slide scanning and machine learning. Stained tissue sections were imaged on a Zeiss Axioscan.Z1 (Carl Zeiss AG, Oberkochen, Germany) at 20x using fluorescence filters configured for DAPI, FITC, and Cy3. Whole-slide .czi files were imported into TissueStudio 2.4 (Definiens AG, Munich, Germany) for automated tissue detection followed by manual correction of ROIs to delineate endometriosis lesion, peritoneum, haemorrhage, and adipose tissue. Tissue studio’s built-in nuclear segmentation, using the DAPI channel, was applied within these regions to identify cell objects and these objects were then classified as positive or negative for each channel based on intensity thresholds which were used across all samples.

### Statistical analysis

Statistical analysis was carried out in GraphPad Prism 7.02. Data was first analysed for normality using an Anderson Darling normality test. If data were normally distributed, either an ANOVA with a Tukey’s post-hoc test (more than 2 samples) or a t-test (2 samples) was performed. If data were not normally distributed, non-parametric tests were used; either Kruskal-Wallis with a Dunn’s post hoc test (more than 2 samples) or a Mann-Whitney U test (2 samples). Statistical significance was reported at p<0.05.

## Acknowledgements

We thank the QMRI flow cytometry and cell sorting facility technicians (University of Edinburgh) for advice on panel design, and Fiona Ballantyne for technical assistance. We thank Ronnie Grant (University of Edinburgh) for help with figure preparation. This work was supported by a Medical Research Council (MRC) Career Development Award (MR/M009238/1; to E.G), a MRC Project Grant (MR/S002456/2; E.G), an MRC Centre studentship to C.H, a Wellcome Trust Senior Investigator Award (101067/Z/12/Z; J.W.P) and an MRC Centre Grant (MR/N022556/1; J.W.P). The authors declare no conflict of interest.

## Author contributions

E.G conceived experiments, analysed data and wrote manuscript; C.H conceived and performed experiments and analysed data. P.D and M.R performed experiments. M.M provided reagents and critical feedback. D.S performed image analysis; J.P provided critical discussion and advice on manuscript. S.J provided advice on experimental design, interpretation, critical feedback and manuscript preparation. A.W.H provided feedback on experimental design and manuscript preparation.

