## Supplementary Figures for "Macrophages inhibit and enhance endometriosis depending on their origin"

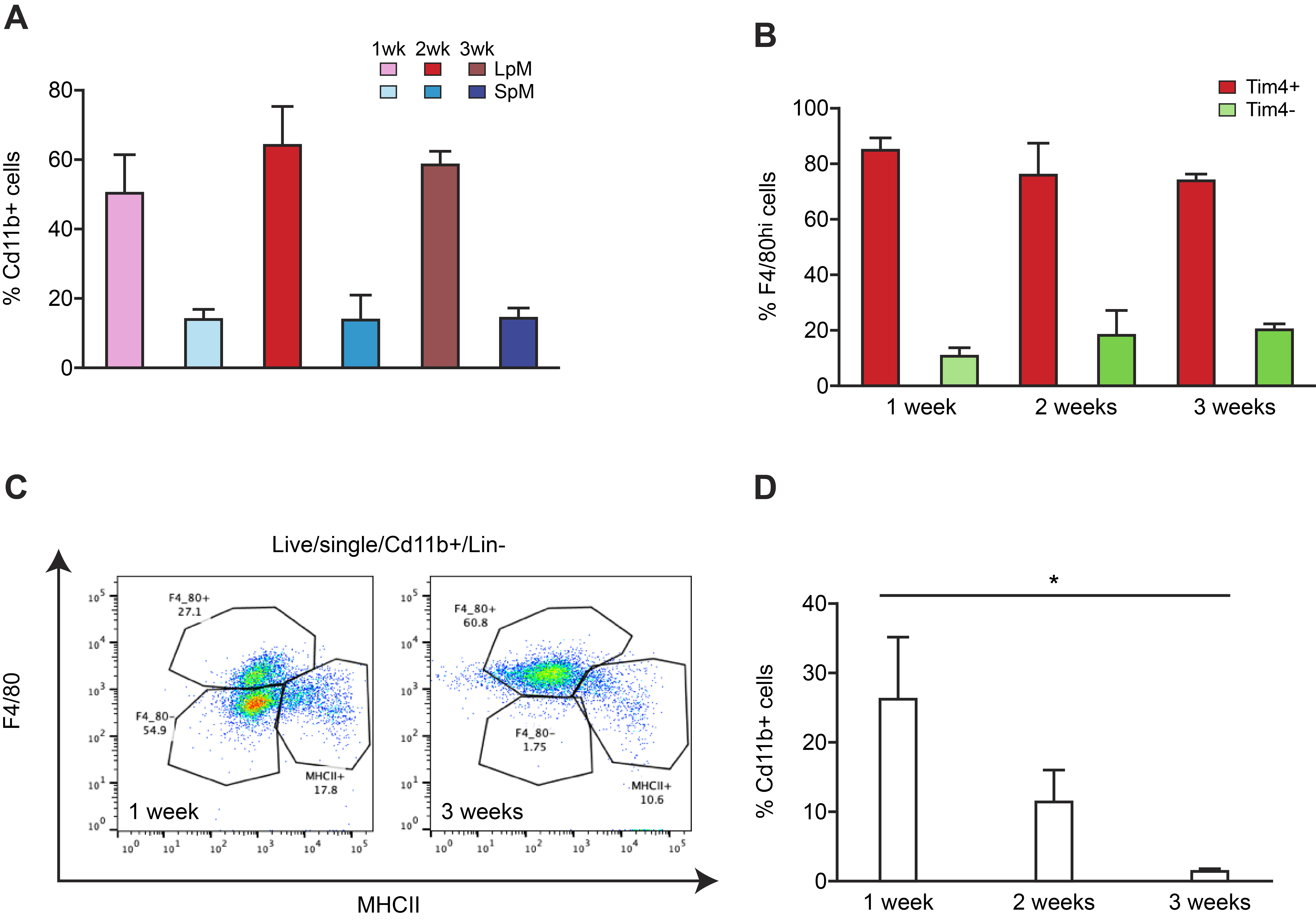

**Figure S1: Large and small peritoneal macrophage ratios are not perturbed in a minimally invasive mouse model of endometriosis.**

A) Endometriosis was induced in recipient mice that had not had their ovaries removed (n=10). LpM and SpM were quantified by flow cytometry at 1 week (n=4), 2 weeks (n=3), and 3 weeks (n=3) post tissue injection. Lesions were recovered from all mice.

B) Quantification of TIM4+ and TIM4- cells in the F4/80^hi^ LpM population present in the peritoneal lavage fluid of mice with endometriosis.

C) LpM and SpM populations in the peritoneal lavage fluid of mice with induced endometriosis 1 and 3 weeks after endometrial tissue injection.

D) Quantification of the F4/80^lo^, MHCII^lo^ population in peritoneal lavage fluid.

Statistical significance was determined using a Kruskall-Wallis test. *;p<0.05.

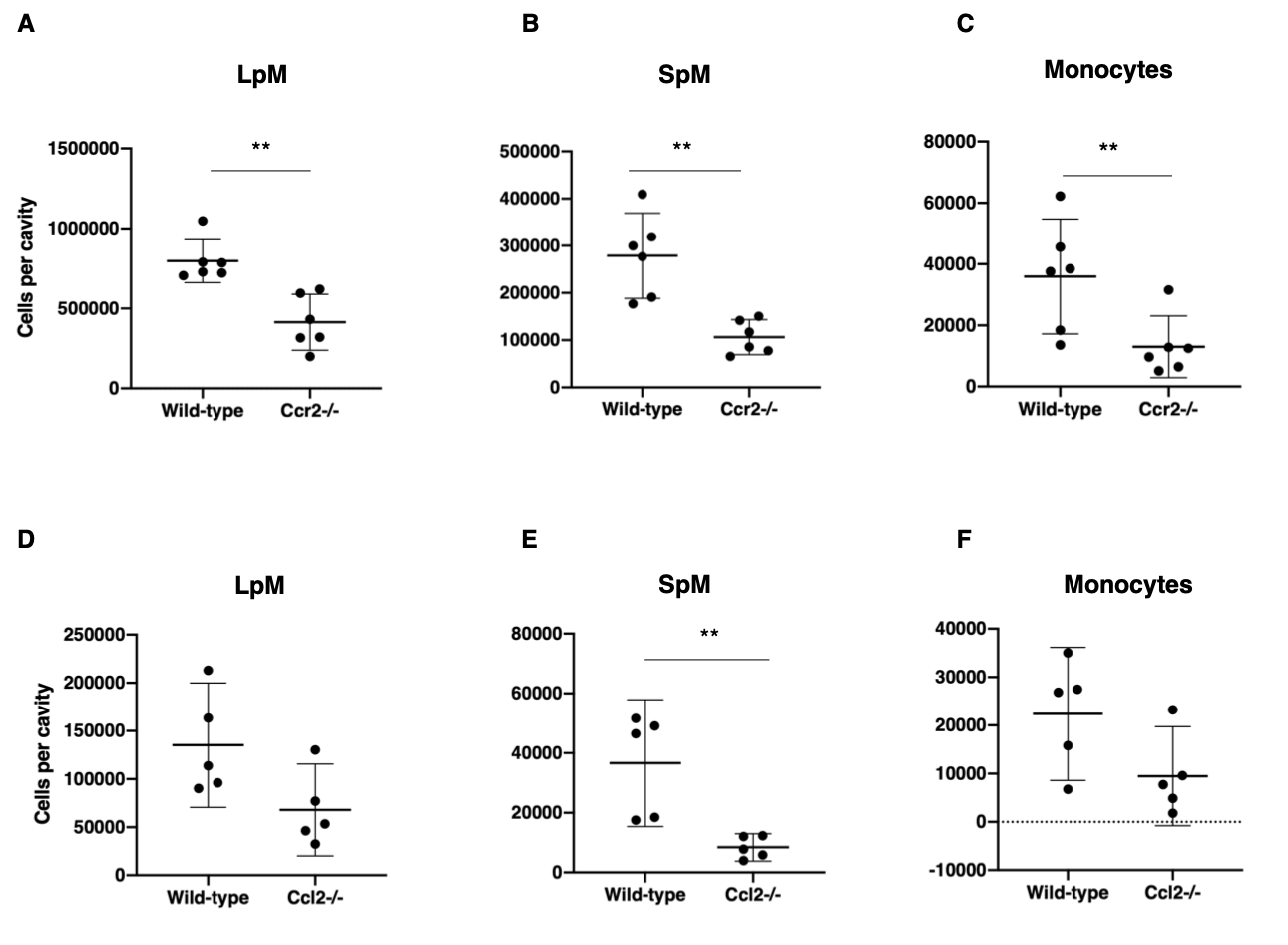

**Figure S2: Absolute numbers of LpM, SpM and monocytes in wild-type vs *Ccr2*-/- or *Ccl2*-/- mice with induced endometriosis.** A-C) Quantification of LpM (A), SpM (B), and monocytes (C) per cavity in wild-type vs *Ccr*-/- mice with induced endometriosis. D-F) Quantification of LpM (D), SpM (E), and monocytes (F) per cavity in wild-type vs *Ccl2*-/- mice with induced endometriosis. Cells per cavity were calculated using count beads during cytofluorimetric analysis. Data represented are mean with 95% confidence intervals. Statistical significance was determined using a Mann-Whitney test. **;p<0.01.

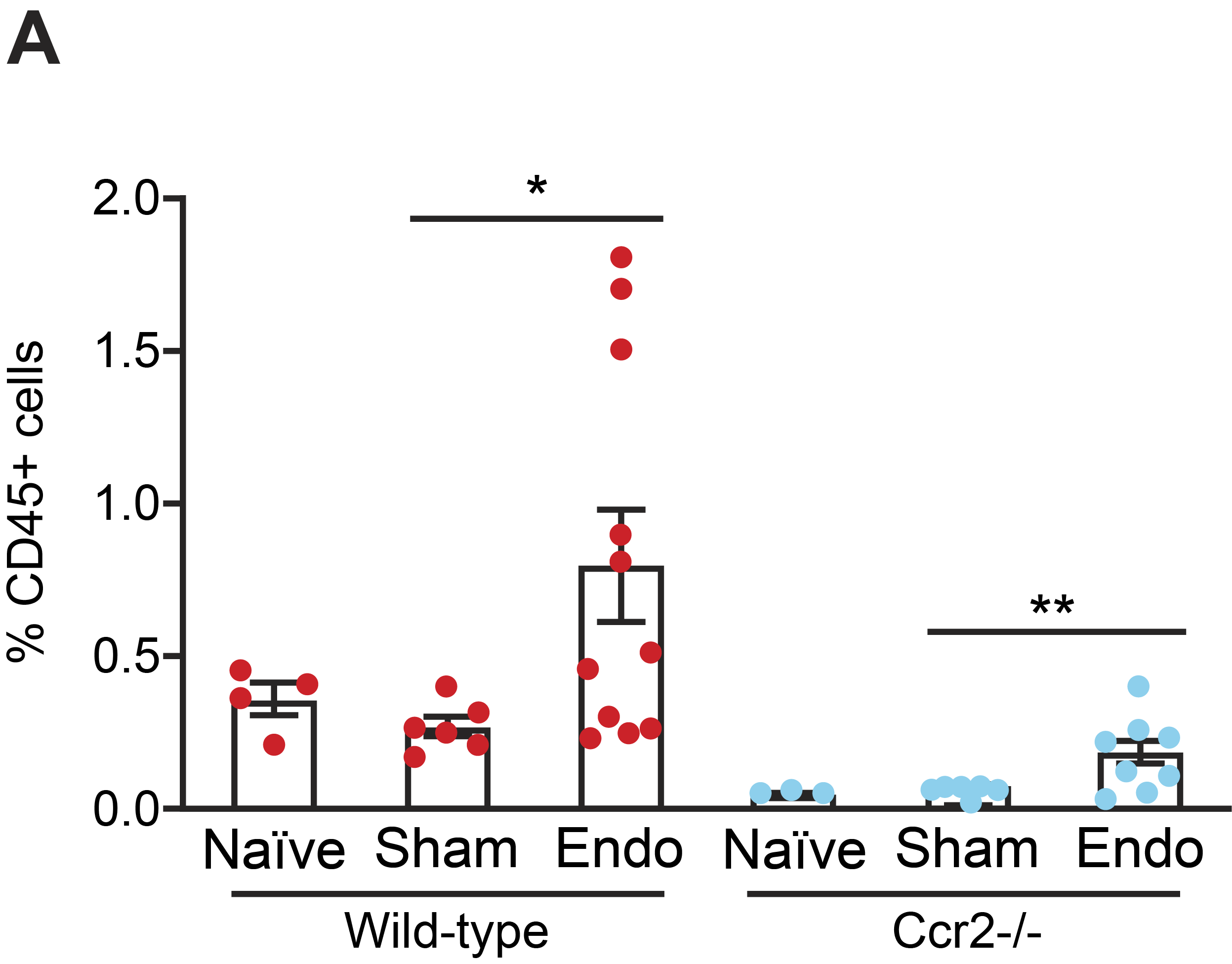

**Figure S3: Monocytes can be recruited to the peritoneal cavity of Ccr-/- mice.**

A) Quantification of monocytes (Ly6C^hi^ cells) in naïve, sham and mice with induced endometriosis (wild-type vs *Ccr2*-/- mice).

Data are presented as mean ± SEM. Statistical significance was determined using a one-way ANOVA and a Tukey post-hoc test, *;p<0.05, **;p<0.01.
